# DIFFERENTIAL REGULATION OF GENES IN ENTORHINAL CORTEX AND HIPPOCAMPUS IN LATE ONSET AD

**DOI:** 10.1101/2024.09.14.613003

**Authors:** Moumita Biswas, Soumalee Basu

## Abstract

Amyloidogenic processing of APP mediated by increased BACE1 activity resulting in Aβ production and accumulation of hyperphosphorylated tau cause neuronal degeneration in AD. Pathological changes occur early in the EC and the hippocampus, much before neurodegeneration sets in. Here, using 500 and 303 differentially expressed genes from the EC and hippocampus respectively, we observed impairment of several cellular functions owing to either disrupted transcription or post-translational regulation, through functional enrichment, PPI network, miRNA and convergent functional genomic analysis. In EC, functional enrichment analysis highlighted several processes of which the Oncostatin-M-mediated pathway is related in the disease pathogenesis by increasing BACE1 transcription through STAT3 activation. Hub genes, BRD4 and PIN1, involved in Aβ and tau pathology are downregulated probably through the concerted action of hsa-miR-3613-3p and hsa-miR-4674, upregulated miRNAs in AD. In the hippocampus, upregulated FXR1 and downregulated RTN3 being involved in post-transcriptional processing of BACE1, result in increased Aβ production. For the disruption observed in the autophagy-lysosomal pathway, neurotransmitter synthesis and Aβ clearance, YWHAH, a downregulated hub gene is instrumental. Regulation of YWHAH expression is probably mediated at the post-transcriptional level through hsa-miR-455-3p and hsa-miR-4674, known to be heavily upregulated in AD. In both the regions, similar pattern of disrupted cellular pathways emerge, although the key players, differ. Whereas dysfunctional proteasomal degradation is mediated by SKP1-CUL1 in EC, it is CUL3 mediated in hippocampus. Thus, similar disease pathology develops through dysregulation of pathways that are evidently different in EC and hippocampus implying failure of therapeutics based on single target.

## 1.0 Introduction

Neurons of the unbelievably complex human nervous system, not only form intricate connections with each other but are also connected with glial cells, astrocytes, oligodendrocytes and so on to form a processing unit that is structurally and functionally different from the rest of the body (Sousa *et al*. 2017). Each distinct region of the brain is involved in a particular function covering a plethora of activities, most of which are affected for a person afflicted with Alzheimer’s disease (AD). In phases, these patients lose the ability to perform specialised tasks that in any normal person remain unaffected throughout the entire lifespan. Beginning with minor forgetfulness and later developing into full-fledged dementia (Ballard *et al*. 2011; Masters *et al*. 2015), the disease is becoming a major healthcare challenge worldwide.

Brain imaging studies indicate that medial temporal lobe atrophy of the entorhinal cortex (EC) at a preclinical stage and of the anatomically related structure hippocampus at a later stage is a pathophysiological event characterising AD brains macroscopically (Coupé *et al*. 2019). AD brains are also characterised by the presence of prototypical lesions like extracellular protein plaques and intracellular neurofibrillary tangles. The chief constituent of the extracellular plaques are aggregates of the insoluble amyloid β peptide formed by the sequential processing of the amyloid precursor protein (APP) by β-secretase (BACE1 enzyme) and γ-secretase respectively. The intracellular neurofibrillary tangles are formed from the hyperphosphorylated microtubule associated tau protein (Sathya *et al*. 2012; Liu *et al*. 2019; Lopez *et al*. 2019).

Decades of research carried out to shed light on the pathways initiating the onset of the disease have shown limited success with the exact cause of the disease still remaining elusive (Breijyeh *et al*. 2020). Also, the mechanisms by which these changes in the brain lead to cognitive decline are debated till date. Anatomical studies from postmortem AD patients revealed that abnormal changes very likely start from the EC (Van Hoesen *et al*. 1991).Moreover, in all cases of Alzheimer’s disease, EC have been a consistent focus of pathology with selective changes that altered some layers more than others. Interestingly, neurons of the EC receive input from multiple cortical regions and send axons to the hippocampus suggesting that an early dysfunction in the activity of EC marked by mild forgetfulness gradually affects the hippocampus which is manifested as dementia at a later stage. At this juncture, it was thought that analyses of differentially expressed genes of specific brain regions in response to pathological conditions could be fruitful in identifying key molecular pathways associated. The aim of this work was identification and comparison of differential gene expression profiles in the entorhinal cortex with hippocampus of late onset AD patients.

In this study, gene expression profiles from post-mortem brains of AD patients were acquired from the GEO database and using a software package, a differential gene expression analysis was carried out to explore change in gene expression between healthy and diseased conditions. Functional annotations were enriched using two approaches. Thereafter, resulting DEGs were integrated with protein–protein interaction (PPI) network. Clustering module analysis and centrality analysis followed by convergent functional genomic analysis were performed to prioritise and rank the candidate hub genes. This knowledge of the relevant and distinct pathways and their interactions, if any, may be useful for the prevention of AD pathology or delaying the onset of dementia.

## 2.0 Materials and methods

### 2.1. Data retrieval and differential gene expression (DEG) analysis

Microarray dataset GSE5281 on Affymetrix Human Genome U133 Plus 2.0 Array platform [HG-U133_Plus_2] was downloaded from the NCBI repository GEO DataSets (Liang *et al*. 2008; Readhead *et al*. 2018). The dataset contains 161 post-mortem brain samples of AD patients and non-demented normal control (CN) subjects collected from three Alzheimer’s disease centres corresponding to six regions namely entorhinal cortex, hippocampus, medial temporal gyrus, posterior cingulate, superior frontal gyrus, primary visual cortex and middle temporal gyrus of which only the files with extension .cel was retrieved from the entorhinal cortex region data for analysis.

The pre-processing of the datasets and their subsequent DEG analysis was done using various libraries of the R software package BiocManager. R package Affy that uses the robust multi-array normalisation technique and quantile normalisation was used to createthe expression matrix (Gautier *et al*. 2004). Then the expression dataset was duly annotated and processed through the Limma package to identify the differentially expressed genes (DEGs) (Ritchie *et al*. 2015).

### 2.2. Gene set enrichment analysis

Gene ontology (GO) enrichment analyses were performed in R using the function of clusterProfiler (Yu *et al*. 2012; Wu *et al*. 2021). The R package EnrichR was used to perform the KEGG pathway analysis (Chen *et al*. 2013). Functional and pathway enrichment analyses were conducted separately for upregulated and downregulated genes. In this analysis, a p-value < 0.05 was considered significant for the screening of significant GO terms and KEGG pathways.

### 2.3. PPI network construction

To further explore the interactions among DEGs, PPI network was constructed in the Cytoscape using experimental interaction data downloaded from the Biogrid database (BIOGRID-MV-Physical-4.4.233.tab3) (Shannon *et al*. 2003). In doing so the interaction type was limited to “human”, next log FC was used as a filter. It is to be mentioned that the nodes directly linked with the DEGs were also included.

### 2.4. Hub gene identification using MCODE and centrality analysis

To determine the potential hub genes of AD, Cytoscape pluggin MCODE was used in default settings to identify important clusters (Bader *et al*. 2003). CytoHubba plugin was used to calculate the score for each node (Chin *et al*. 2014). This tool calculates eleven centralities from the provided network out of which the maximal clique centrality (MCC) and bottleneck centrality (BN) was used to show the top 25 ranked nodes.

### 2.5. Upstream regulation study

Hub genes identified from MCODE clustering analysis and centrality analysis were selected as query targets for subsequent miRNA regulation analysis for studying their upstream regulation. The TargetScan database was utilised for extracting information on the miRNAs that bind to the 3’UTR regions of target hub genes (McGeary *et al*. 2019). The resulting data, compiled in Excel sheets, was imported into the R software for the identification of miRNAs that regulate the largest number of these hub genes, offering insights into potential regulatory mechanisms within these brain regions.

### 2.5. Convergent Functional Genomic Analysis

Convergent functional genomic (CFG) analysis integrates biological evidence from multiple sources to prioritise candidate genes implicated in complex diseases. In the study, the Alzdata database, which integrates four evidences namely, genetic association of DNA variation with disease susceptibility, gene expression regulated by AD genetic variants, protein-protein interaction with AD core proteins, and diagnosis prediction of disease models, have been used to conduct a CFG analysis to rank the hub genes and identify those that are differentially expressed at the early stage of the disease before the appearance of AD pathology (Xu *et al*.2018).

## 3.0 Results

### 3.1. Sample characteristics

Gene expression data of 10 samples from patients with AD and 13 samples from age matched control subjects (CN) were analysed using the Microarray dataset GSE5281 on Affymetrix Human Genome U133 Plus 2.0 Array platform. Detailed list of sample characteristics for the entorhinal cortex and hippocampus is provided in **Table 1**.

**Table 1:** Sample characteristics of GSE5281 entorhinal cortex (left panel) and hippocampus (right panel)

| Attributes | Entorhinal Cortex |  | Hippocampus |  |
| --- | --- | --- | --- | --- |
| Age<br>(years) | AD | CT | AD | CT |
| | $84 \pm 5.5$ | $80 \pm 9.2$ | $77.8 \pm 5.71$ | $79.61 \pm 9.40$ |
| No. of samples | 10 | 13 | 10 | 13 |
| RNA sample | 10 ug |  | 10ug |  |

### 3.2. Identification of DEGs in the entorhinal cortex and hippocampus

Recently, brain imaging studies from preclinical AD patients and electrophysiological recordings from AD animal models have shown that impaired neuronal activity in the EC precedes neurodegeneration (Harris *et al*. 2010; Khan *et al*. 2014). We attempted to analyse changes in gene expression confined to this region of the brain in AD patients. After background correction and normalisation, gene expression of 54,645 genes was assessed. Analysis with threshold set at adj.P.Val< 0.001 and log FC > 1.4 and < −1.4 was carried out for both the brain regions. In the entorhinal cortex a total of 500 genes were found to be differentially expressed with 109 upregulated (21.8%) and more than a double, i.e.391 downregulated genes (78.2%). A volcano plot showing the significantly upregulated and downregulated genes is shown in **Fig.1.(A)** *FAM107B, TJP2, HIPK2, SPP1,* and *ERBB2IP* are the top 5 upregulated genes while *MIF*, *CHRM1*, *STYK1*, *RBP4*, and *AP2M1* are the top 5 down-regulated genes (**Table S1 & S2**).Out of the top five upregulated genes, two have no known biological function, while from the remaining three only *TJP2* probably has disease mitigating role in AD owing to its involvement in blood brain barrier restoration (Fanning *et al*. 2009). The remaining two genes, *SPP1* and *HIPK2* probably have roles in the disease pathogenesis (De Schepper *et al*. 2023; D’Orazi *et al*. 2012). Upregulated SPP1 might result in dysfunctional microglia-mediated synaptic phagocytosis and/or phagocytic activity of reactive astrocytes thereby severely affecting functional neuronal communication. HIPK2 upregulation could be implicated in ER stress-mediated neurodegeneration. On the other hand, the top 5 downregulated genes are involved in distinct biological processes. *MIF* is a proinflammatory cytokine while loss of *CHRM1* results in mitochondrial dysfunction and impairment of respirasome assembly, *STYK1* is known to be involved in macroautophagy, *RBP4* transports retinol in association with transthyretin, and *AP2M1* is involved in sorting and endocytosis (Nasiri *et al*. 2020; Dean and Scarr, 2021; Zhou *et al*. 2020; Ishi *et al*. 2019; Tian *et al*. 2013). Dysregulated synaptic pruning, poorly regulated energy metabolism, compromised blood brain barrier and chronic pro-inflammatory response emerge as highlighting features characterized by the top up- and downregulated genes of EC.

**Fig.1:**
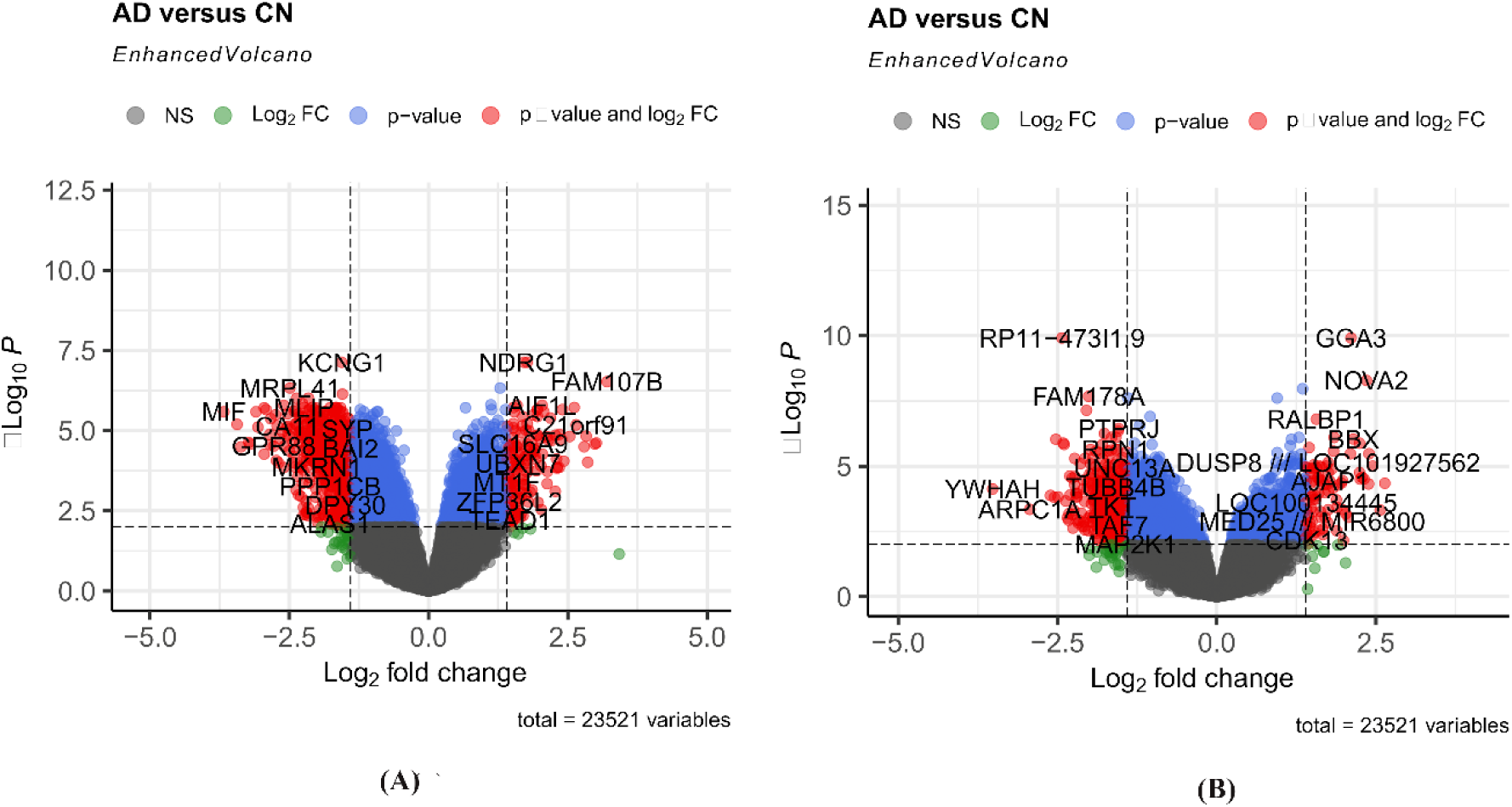
Volcano plot showing the significant differentially expressed genes of AD versus cognitively normal (CN) samples from (A) entorhinal cortex (B) hippocampus.

In the hippocampus, out of the 303 genes that are differentially expressed, 247 (81.5%) genes are downregulated and 56 (18.4%) genes are upregulated **(Table S3 & S4)**. LOC202181, FXR1, MINK1, SRRM2, and NOVA2 are the top 5 upregulated genes while YWHAH, ARPC1A, CASD1, CD9 and RTN3 are the top 5 down-regulated genes (**Fig.1. (B)**). Interestingly, none of the top genes differentially expressedin the EC are associated with amyloidogenic processing of APP, but in the hippocampus two such genes have been found; one - RTN3 or reticulon 3, heavily downregulated, that is exclusively expressed in the neurons and negatively regulates β-secretase activity of BACE1 (He *et al*., 2004; Sharoar and Yan, 2017); two - FRX1, highly upregulated, and binds to the 5’ UTR of BACE1 increasing its translation (Zhou *et al*. 2023). Broadly speaking, among the remaining upregulated genes, NOVA2 (Ule *et al*. 2005) and SRRM2 (Xu *et al*. 2022) function in alternative splicing of mRNA and MINK1 is involved in the trafficking of AMPA receptors at the surface of hippocampal neurons. It has been shown that ER stress driven phosphorylation of SRRM2 results in its mislocalisation from nuclear speckles tocytoplasm wherein pathological tau promotes pSRRM2 accumulation even in the absence ofAβ. Thus it implies that DEGs of the hippocampus are not only associated with amyloidogenic processing of APP but also associate with tau pathology. Within the downregulated genes, YWHAH is important for the regulation of the autophagy-lysosomal pathway (ALP) as well as for the synthesis of neurotransmitters dopamine, adrenaline and serotonin (Bruet *et al*. 2016), CASD1 is a key enzyme in the biosynthesis of 9-O-acylated sialoglycans (Baumann *et al*. 2015), and CD9 is a general anti-inflammatory marker of monocytes and macrophages with diverse biological role (Umeda *et al*. 2020). From our findings, we can infer that the features of AD brain as proposed by the amyloid cascade hypothesis and/or tauopathymay be explained better from the DEGs in the hippocampus than in the entorhinal cortex. It is to be remembered that in the EC, neural circuits are refined by selecting neurons and neural pathways through synaptic pruning depending on their levels of activity. SRRM2 as DEG might impair the regulation of this refinement leading to the early space time disorientation of AD patients (Tanaka *et al*. 2018). Since entorhinal cortex is perceived as an input and output structure of hippocampus, neuroinflammation and or neurodegeneration in EC would resultin hippocampal dysfunction affecting learning and memory.A more direct involvement of memory impairment in hippocampus could be throughupregulated MINK1, disrupting regulation of AMPA receptor trafficking to synapses and affecting long-term potentiation.However,why hippocampus forms the seat of tauopathy and increased amyloidogenic processing of APP remains obscure.

The genes listed in **Table S1 & S2** and **Table S3 & S4** are ranked by order of their fold change, but it is important to note that these top-ranking genes may not necessarily represent the critical changes that occur in the early stages of late onset AD. To gain a better understanding of the potential key regulators of the disease, additional investigations were carried out. These involved functional enrichment, protein-protein interaction network analysis, centralityanalysis andidentification of micro-RNAs in the posttranscriptional regulation of hub genes, all aimed at identifying potential key players of the disease.

### 3.3. Functional annotation using gene ontology and KEGG pathway analysis

Gene ontology (GO) enrichment analysis was performed on the dataset to elucidate their biological functions in the entorhinal cortex and hippocampus of AD patients as shown in **Fig.2**. In the entorhinal cortex (**Fig.2(A)**), for the category of biological processes, upregulated genes that are involved in the Oncostatin-M-mediated pathway (GO: 0038165), stress response to copper ion (GO: 0046688), and detoxification of copper ion (GO:0010273) are significantly enriched while down-regulated genes are significantly enriched in the mitochondrial translation (GO:0032543), mitochondrial gene expression (GO: 0140053), and aerobic electron transport chain (GO: 1990169). Of particular interest, the upregulated Oncostatin-M mediated pathway may be associated with an increased amyloidogenic processing of APP since its downstream effector STAT3 is a transcription factor and positive regulator of BACE1 activity (Millot *et al*. 2020). Similar trend is also observedin hippocampus (**Fig.2(D)**). In both the regions, pathways related to oxidative phosphorylation and ATP production in mitochondria are enriched among the downregulated DEGs. However subtle differences can be observed for the upregulated genes. While pathways that are activated in response to accumulation of metal ions, in particular copper ions and Oncostatin-M-mediated pathway are enriched in the entorhinal cortex, in the hippocampal region we observed enrichment of nitric oxide signalling pathways along with transmembrane polyol transport and mechanosensory behaviour.

**Fig. 2:**
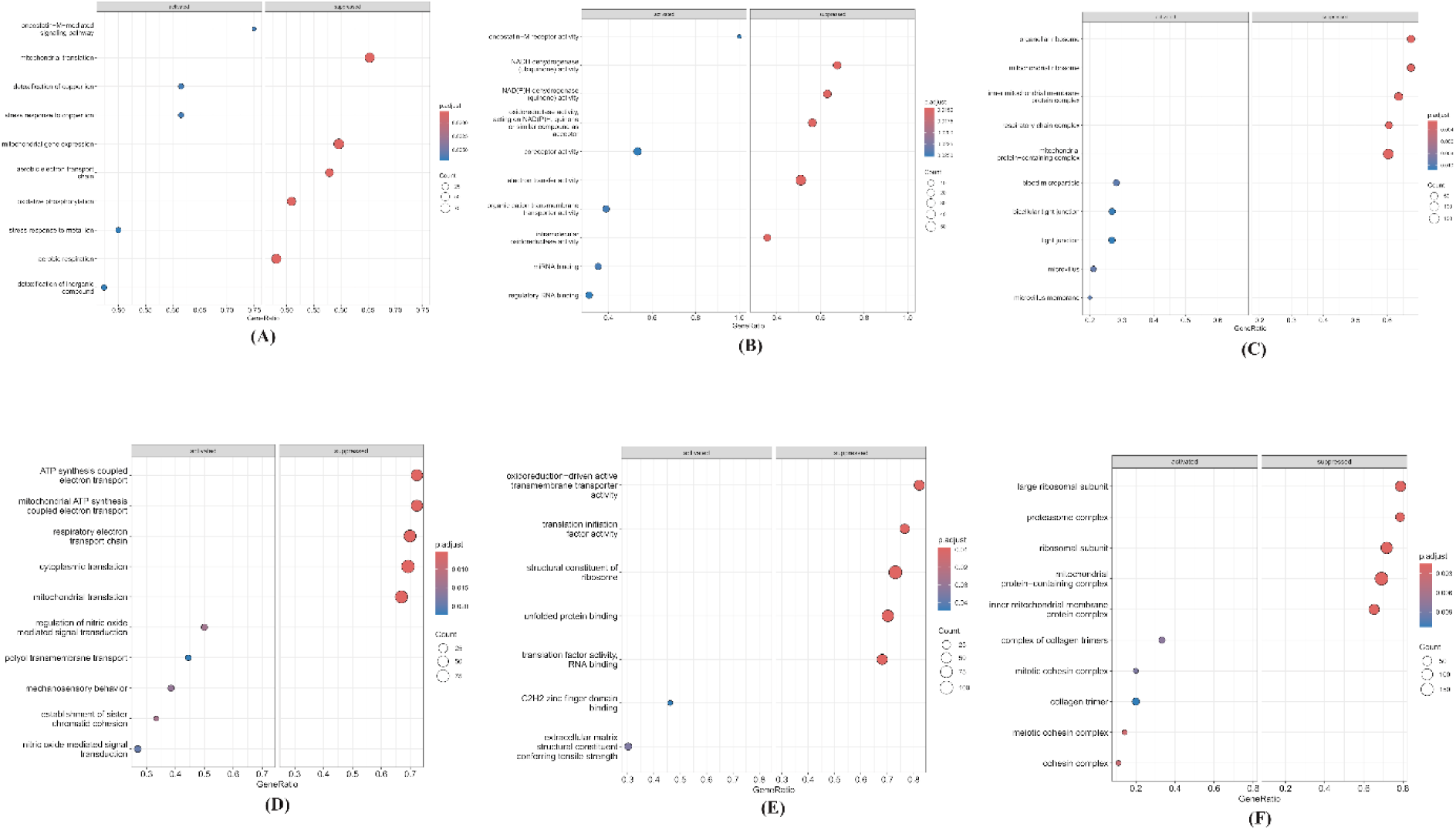
Gene Ontology of top (A) biological processes, (B) molecular function and (C) cellular component of entorhinal cortex (top panel) and (D) biological processes, (E) molecular function and (F) cellular component of hippocampal region (bottom panel)

In the molecular function category of EC (**Fig.2(B)**), upregulated genes were enriched in the Oncostatin-M receptor activity (GO: 0004924), coreceptor activity (GO:00150256), organic cation transmembrane transporter activity (GO: 0015101), while the downregulated genes were enriched in the NAD(P)H dehydrogenase (quinone) activity (GO: 0003955), oxidoreductase activity acting on NAD(P)H quinoneor other similar compounds (GO: 0016491), NADH dehydrogenase (ubiquinone) activity (GO: 0008137). For the cellular components category (**Fig 2(C)**), upregulated DEGs were found to be enriched in blood microparticle (GO: 0072562), tight junctions (GO: 0070160), and bicellular tight junctions (GO: 0005923) while the downregulated genes were found to be enriched in organellar ribosome (GO: 0000313), mitochondrial ribosome (GO: 0005761), and inner mitochondrial membrane complex (GO: 0005923). In the molecular function (**Fig.2(E)**) and cellular component category (**Fig.2(F)**) no noteworthy differences in enriched pathways were observed between the entorhinal cortex and hippocampus region. Enrichment analysis with the upregulated genes show enrichment of specific pathways, while for the downregulated genes (more in number than upregulated genes) a number of pathways primarily associating the bioenergetic processes are enriched. Biological process, molecular function and cellular component all point towards mitochondrial dysfunction. However, the dysfunction in the two regions shows dysregulation at different levels. In EC, the expression of the electron transport chain (ETC) components and that of the mitochondrial ribosomal subunits are low signifying dysregulation both at the transcriptional level of the nuclear encoded genes and at the translational level of the mitochondrial encoded genes. In the hippocampus however the dysregulation is limited to the transcriptional level with majority of the differentially expressed genes corresponding to a different set of the ETC components. This implies dysregulation of translation (repression) of the mitochondrial encoded ETC proteins thus supporting the energy crisis in Alzheimer’s disease.

KEGG analysis is used to interpret the functional significance of large sets of genes or proteins. Following a GO analysis, a KEGG pathway enrichment analysis was performed for both the regions (**Fig. 3**) in order to further evaluate the biological roles of the DEGs.According to the KEGG analysis, pathways involved in Parkinson’s disease (p.value: 2.69×10^−5^) and Huntington’s disease (p.value: 3.11×10^−5^) were highly enriched in both the regions.Interestingly, in the entorhinal cortex(**Fig. 3(A)**), the retrograde endocannabinoid pathway emerged as the most significant pathway (p.value: 2.27×10^−5^) and is possibly linked with the disease pathology. The pathway is known to suppress excitatory neurotransmitter release (Lu et al. 2016) the associated genes of which are downregulated. This implies excessive activation of glutamate or other excitatory receptors eventuating in neuronal dysfunction and death. Thus the affected retrograde endocannabinoid signalling pathway could be held responsible for the excitotoxicity owing to excessive activation of the NMDA receptors that is thought to be closely associated with localised neuronal viability in AD pathogenesis (Wang and Reddy, 2017).

**Fig.3:**
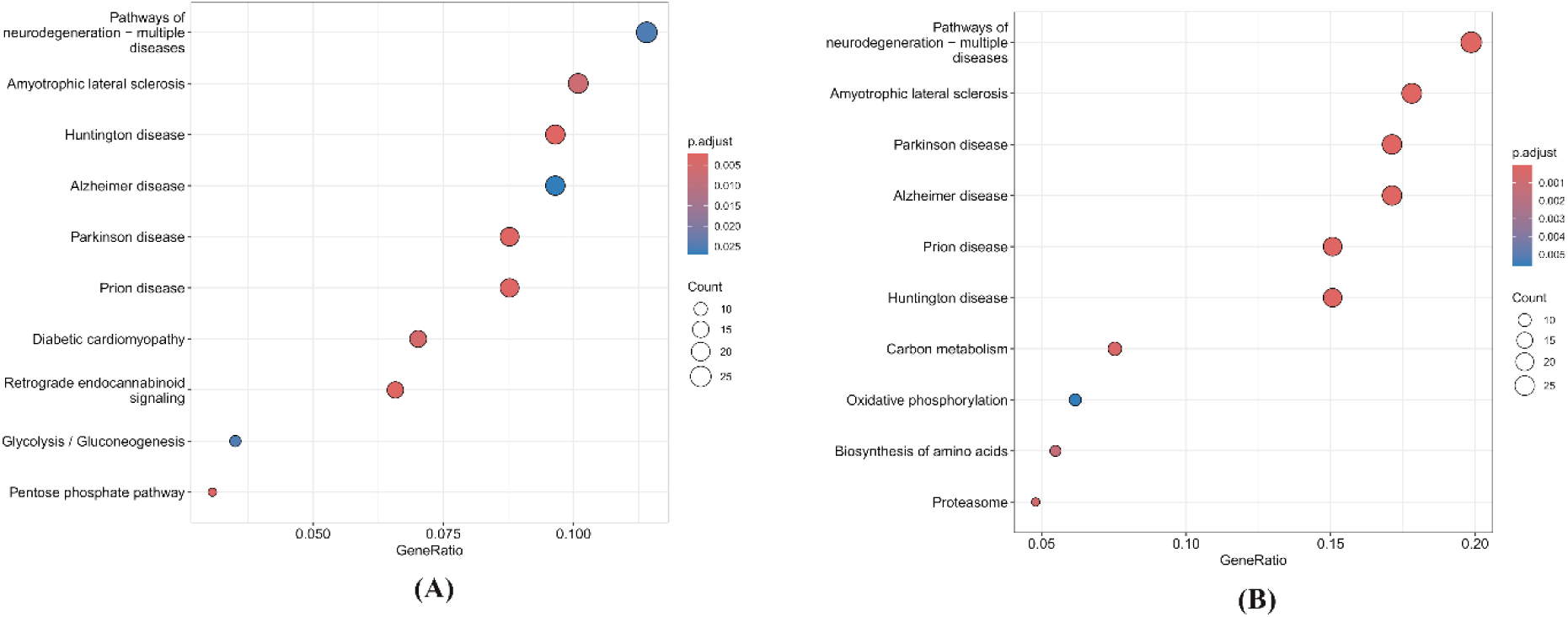
KEGG pathway enrichment analysis of the identified DEGs from the (A) entorhinal cortex and (B) hippocampus region.

### 3.4. Identification of hub genes using PPI network

In the study, the protein-protein interaction network of the entorhinal cortex and hippocampus was created using physical interaction information from the BioGRID database. The PPI network has 3130 nodes with 11170 interactions and 3124 nodes and 14037 interactions for the entorhinal cortex and hippocampus respectively. Although the number of nodes is comparable, the number of interactions in hippocampus is higher than EC.

#### 3.4.1. MCODE clustering analysis

MCODE utilises graph theoretic concepts to detect highly interconnected subgraphs that are likely to represent functional modules or protein complexes. It then assigns scores to the detected complexes based on various characteristics, such as the number of nodes, the density of connections and the size of the complex (Zaki *et al*. 2012). The scoring helps to prioritise and rank the complexes based on their significance. In our study, the three most significant modules cluster I, II and III, selected on the basis of their module score are shown in **Fig.4**.

**Fig. 4:**
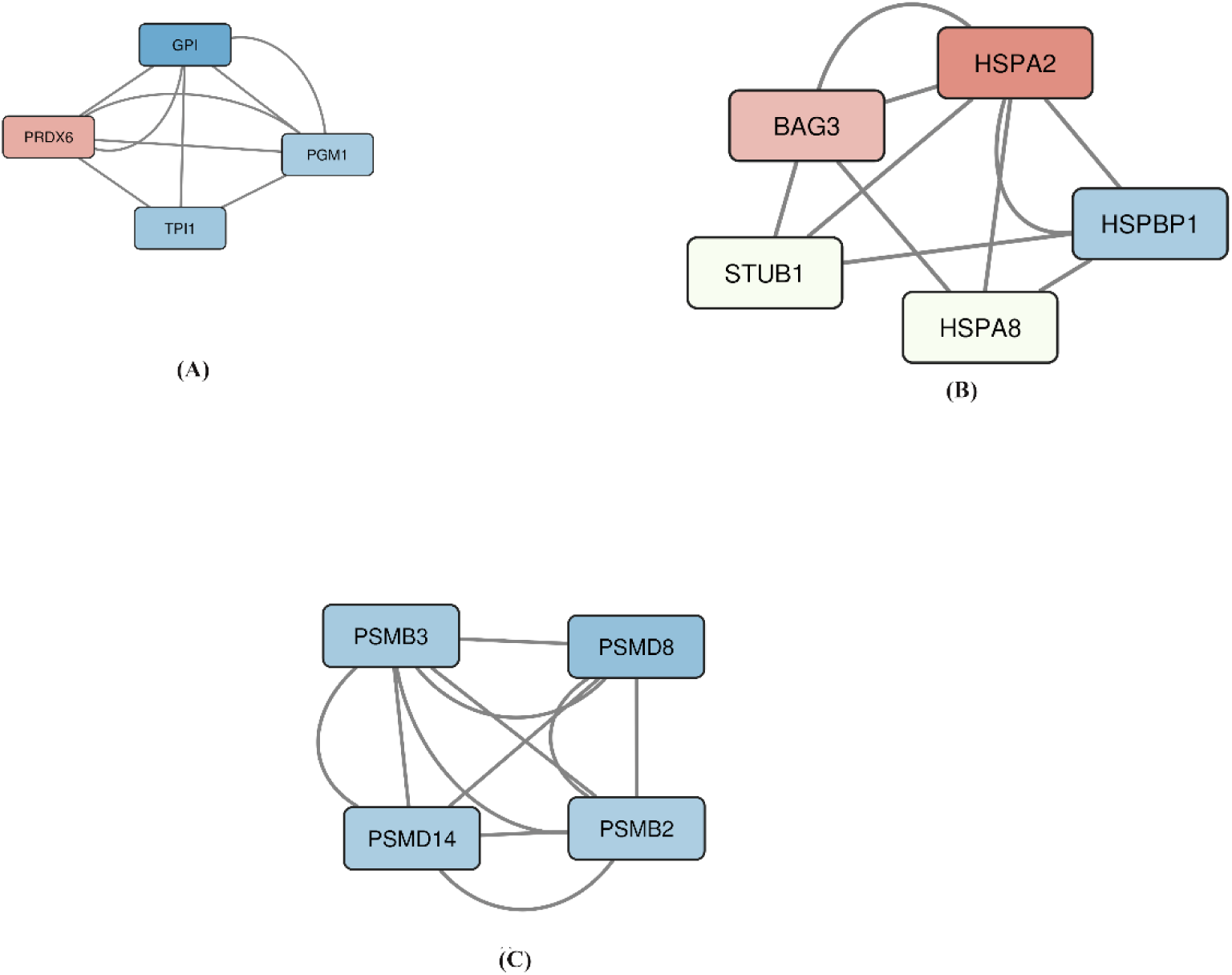
Top three cluster modules of the entorhinal cortex region, (A) cluster I (B) cluster II (C) cluster III using MCODE. Colour representation is based on logFC; the darker shade represents larger logFC. (red: upregulated, blue: downregulated, white: no change in expression)

In the first network, the most significant module of EC, **Fig.4(A)**, consisted of 4 nodes, with 3 downregulated nodes and one upregulated node, namely, GPI, PGM1, TPI1, and PRDX6 respectively. Notably, the three downregulated hub nodes are involved in glucose metabolism, glucose-6-phosphate isomerase (GPI) and triose phosphate isomerase 1 (TPI1) are involved in glycolysis while phosphoglucomutase 1 (PGM1) catalyses the interconversion of glucose-1-phosphate to glucose-6-phosphate in the glycogenolysis pathway. The single upregulated node in the network encodes for peroxiredoxin 6, an antioxidant enzyme probably to mitigate oxidative stress (Arevalo and Vázquez-Medina, 2018) induced lipid peroxidation resulting as an effect of hyperglycemic conditions prevailing due to the downregulated glycolytic enzymes.Transcription factor binding motif analysis revealed the presence of common transcription factors in the upstream of the four nodes of the cluster of which transcription factor SOX 5, THAP 11, IRF4 and GATA4 are an interesting finding since they can act both as a transcriptional activator and repressor and are differentially expressed in our dataset and is possibly responsible for the differential expression of the genes of the module.

The second cluster (**Fig. 4(B)**) has 5 nodes, consisting of two upregulated genes, HSPA2 and BAG3, one downregulated gene, HSPBP1, and two genes (STUB1 and HSPA8) that show no change in expression. With HSPA2 and BAG3 as chaperones, HSPBP1 as inhibitor of STUB1 which is an E3 ubiquitin ligase, the network might be involved in reducing the load of misfolded proteins. The third cluster, (**Fig. 4(C)**), has four nodes namely PSMB3, PSMB2, PSMD8 and PSMD14. All the downregulated nodes are subunits of the core and regulatory subunit of the 20S and 19S units of the 26S proteasomal assembly respectivelythat participate in the degradation of misfolded proteins. PSMD14 has an additional function of transporting macroautophagy protein ATG9 from the Golgi to phagophore in the initial stage of the phagosome assembly (Bustamante *et al*. 2020). The analysis brings out the three important affected pathways namely glucose metabolism, chaperone mediated protein folding and proteasomal degradation as the top three clusters. Although these pathways are already implicated in AD, the differential expression of which specific genes from which brain region led to the condition are key findings of the analysis. Interestingly, the two most significant modules from the hippocampal region (**Fig. 5 (A, B)**) highlighted the proteasomal degradation pathway while the third module (**Fig. 5 (C)**) was a pathway that assists chaperone mediated protein folding.

**Fig.5:**
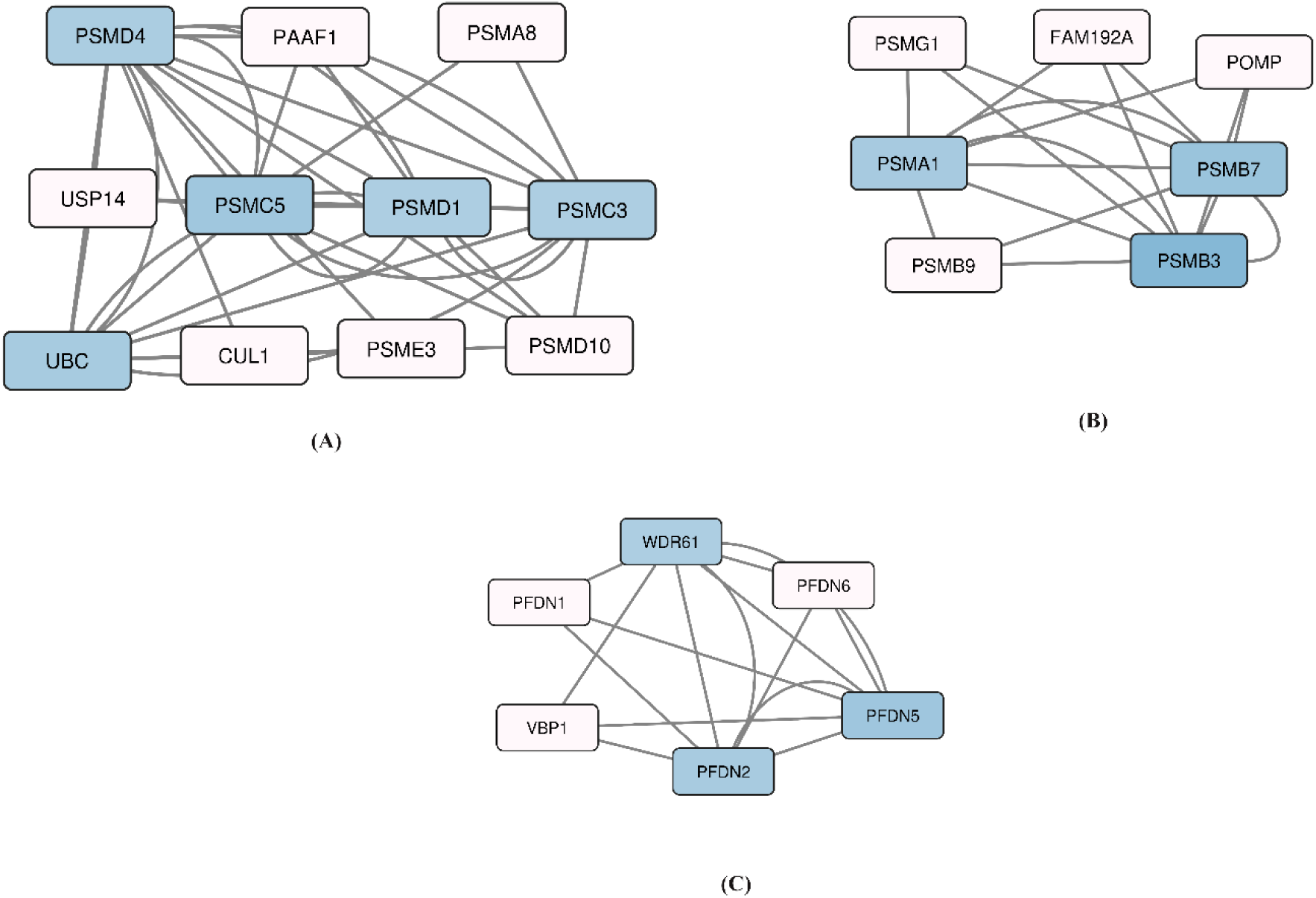
Top three cluster modules of the hippocampus region, (A) cluster I (B) cluster II (C) cluster III using MCODE. Colour representation is based on logFC; the darkest shade represents larger logFC. (red: upregulated, blue: downregulated, white: no change in expression)

#### 3.4.2. Centrality analysis

Centrality analysis has been used in this study to identify hub genes that potentially play a role in the pathogenesis and progression of the disease. Two network centrality algorithms, maximal clique centrality (MCC) and bottleneck centrality (BN), were used to identify central or hub genes from the protein-protein interaction network using the Cytoscape software. MCC assumes that the most critical network elements tend to cluster together, while BN identifies nodes connected through the shortest path.

The top 25 hub genes identified from the entorhinal cortex region by each method are listed in **Table 2 and 3**, respectively. The corresponding diagrammatic representation isshown in **Fig.6 (A) & (B)**. Between the two-centrality analysis, 13 hub genes are found to have been commonly identified, namely, BRD4, PSMD14, SKP1, BAG3, USP11, BSG, HGS, KAT2B, OBSL1, PPP1CB, IGF1R, NFE2L2 and KPNA1. BRD4 emerged as a top hub gene in both MCC and BN centrality analyses, alongside SKP1 and ANLN. BRD4, a member of the BET (Bromodomain and Extra-Terminal domain) family, acts as an epigenetic reader and transcriptional regulator for RNA polymerase II-mediated transcription and is highly downregulated (Donati *et al*. 2018). A downregulated epigenetic reader and transcriptional regulator could probably explain the highnumber of DEGS in the entorhinal cortex.SKP1, another downregulated hub gene, is a core component of the SCF (SKP1-Cullin-F-Box) complex. It functions as the adaptor protein binding to CUL1 and recruits various F-box proteins to form the SCF complex. This role is crucial for enabling the polyubiquitination of diverse substrates targeted by the F-box proteins, leading to their degradation by the 26S proteasome. Thus, the downregulated hub gene SKP1, probably is a key player in first step of protein degradation pathway with subsequent steps being affected by the proteasomal subunits as observed in **Section 3.4.1** generating proteostatic stress. Anilin (ANLN), the only upregulated hubgene encodingan actin binding protein, promotes cell division, migration and contractile ring formation during mitosis (Kučera *et al*. 2021). Besides its role in cytokinesis, anilin also participates - one, as a scaffold for principal myelin proteins and septins (Erwig *et al*. 2019), two, maintains the integrity of tight junctions preventing their disassembly through interaction. It may be mentioned that from pathway analysis presented in **Section 3.3**, tight junction and bicellular tight junctions were enriched, TJP2, a tight junction protein has already been found to be one of the top most upregulated genes (**Table S1**) and other related genes like myelin basic protein (MBP), myelin-associated oligodendrocyte basic protein (MOBP) has been identified as DEGs from our analysis. A probable implication for the upregulation of this hub gene could be to mitigate themyelin damage observed in AD. However, Papuć and Rejdakhave reported that the remyelination process is associated with tau hyperphosphorylation exacerbating the disease pathology (Papuć and Rejdak, 2020).

**Fig. 6:**
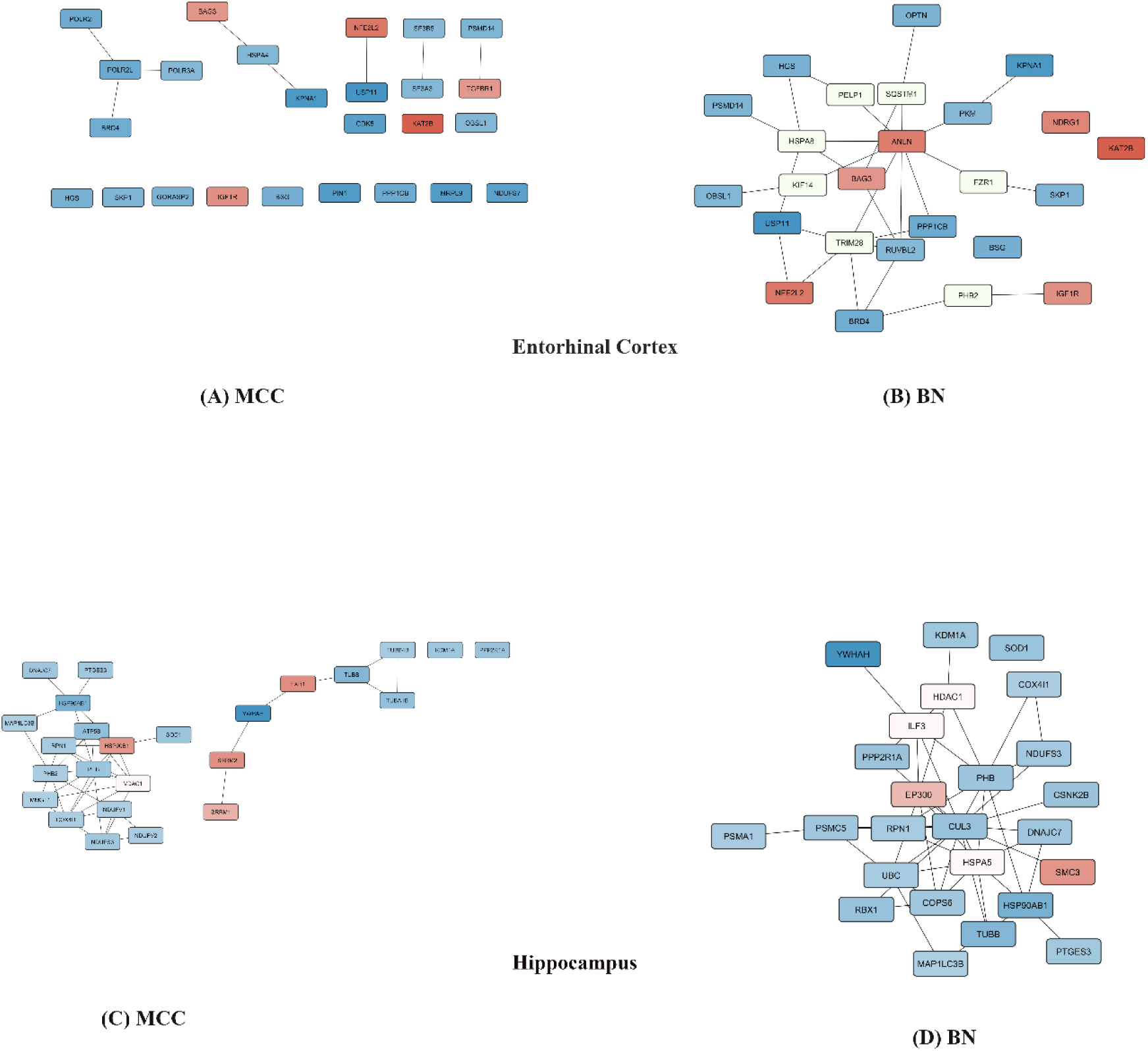
Top 25 hub genes as identified by (A, C) maximal clique centrality (MCC) and (B, D) bottleneck centrality (BN) of the entorhinal cortex (top panel) and hippocampus (bottom panel). Colour representation is based on logFC; the darkest shade represents larger logFC. (red: upregulated, blue: downregulated, white: no change in expression)

**Table 2:** Top 25 hub genes of the entorhinal cortex identified by the maximal clique centrality algorithm analysis.

| **Rank** | **Name** | **Score** |
| --- | --- | --- |
| **1** | **BRD4** | **283** |
| **2** | **SKP1** | **107** |
| **3** | **POLR2L** | **91** |
| **4** | **HSPA4** | **75** |
| **5** | **KPNA1** | **70** |
| **6** | **IGF1R** | **63** |
| **7** | **PIN1** | **61** |
| **8** | **OBSL1** | **59** |
| **9** | **USP11** | **58** |
| **10** | **BAG3** | **55** |
| **10** | **HGS** | **55** |
| **12** | **NFE2L2** | **53** |
| **13** | **MRPL9** | **48** |
| **14** | **POLR3A** | **47** |
| **14** | **PSMD14** | **47** |
| **16** | **PPP1CB** | **45** |
| **17** | **KAT2B** | **44** |
| **18** | **CDK5** | **41** |
| **19** | **BSG** | **40** |
| **20** | **GORASP2** | **39** |
| **21** | **SF3A3** | **37** |
| **21** | **SF3B5** | **37** |
| **23** | **NDUFS7** | **35** |
| **24** | **POLR2I** | **34** |
| **24** | **TGFBR1** | **34** |

**Table 3:** Top 25 hub genes of the entorhinal cortex region identified by the bottleneck centrality algorithm analysis.

| **Rank** | **Name** | **Score** |
| --- | --- | --- |
| **1** | **ANLN** | **1376** |
| **2** | **BRD4** | **1151** |
| **3** | **PSMD14** | **118** |
| **4** | **SKP1** | **117** |
| **5** | **PKM** | **115** |
| **6** | **RUVBL2** | **110** |
| **7** | **HSPA8** | **109** |
| **8** | **BAG3** | **106** |
| **9** | **FZR1** | **91** |
| **10** | **BSG** | **76** |
| **11** | **IGF1R** | **74** |
| **12** | **HGS** | **72** |
| **13** | **USP11** | **70** |
| **14** | **TRIM28** | **68** |
| **15** | **NFE2L2** | **67** |
| **16** | **OBSL1** | **65** |
| **16** | **KPNA1** | **65** |
| **18** | **PHB2** | **64** |
| **19** | **KAT2B** | **63** |
| **20** | **PELP1** | **61** |
| **21** | **SQSTM1** | **60** |
| **21** | **PPP1CB** | **60** |
| **23** | **OPTN** | **59** |
| **24** | **KIF14** | **47** |
| **25** | **NDRG1** | **46** |

Similarly, from the hippocampal dataset, following maximal clique and bottleneck centrality analysis, the top 25 hub genes are tabulated (**Table 4 & *5*)** and the corresponding diagrammatic representation is shown in **Fig.6 (C) & (D)**. 13 genes were found to be common from both the analyses and are YWHAH, COX4I1, PHB, NDUFS3, HSP90AB1, RPN1, KDM1A, TUBB, PPP2R1A, MAP1LC3B, DNAJC7, SOD1 and PTGES3. YWHAH, CUL3 andCOX4I1 came up as top hub genes from both the analyses and are all downregulated. CUL3 is a member of the highly conserved CRL family of proteins that serve a diverse function in the cell, of particular interest is the proteasomal degradation mediated regulation of Nrf2 activity, a key transcriptional regulator of the antioxidant enzymes (Eggler *et al*. 2009). COX4I1 encodes for the largest subunit of the cytochrome c oxidase, component of complex IV of the mitochondrial electron transport chain (Zhang *et al*. 2023). The downregulation of hub genes CUL3 and COX4I1 implicate disrupted bioenergetics and proteasomal degradation that has already been observed from the studies in the previous sections. YWHAH/η (3-monooxygenase/tryptophan 5-monooxygenase activation protein eta), a member of the highly conserved 14-3-3 family of proteins is highly downregulated and is the only hub gene enlisted in the top 5 DEGs.It is required for the synthesis of neurotransmitters dopamine, adrenaline and serotonin as well as in the regulation of ALP. Its downregulation also plays a significant role in clearance of Aβ peptide aggregation since dopamine increases the abundance and activity of the Aβ peptide degrading enzyme, neprilysin (Naoto Watamura *et al*. 2024). It is a striking observation that one of the top DEGs responsible for several biological functions in the hippocampusalso features as a hub gene implying severity of impaired neuronal communication and Aβ clearance leading to Aβ deposition.

**Table 4:** Top 25 genes of the hippocampus region identified from the maximal clique centrality algorithm analysis.

| **Rank** | **Name** | **Score** |
| --- | --- | --- |
| **1** | **YWHAH** | **323** |
| **2** | **COX4I1** | **275** |
| **3** | **PHB** | **261** |
| **4** | **NDUFS3** | **203** |
| **5** | **HSP90AB1** | **127** |
| **6** | **RPN1** | **122** |
| **7** | **KDM1A** | **98** |
| **8** | **PHB2** | **91** |
| **9** | **MMGT1** | **85** |
| **9** | **TUBB** | **85** |
| **11** | **NDUFV1** | **82** |
| **12** | **PPP2R1A** | **80** |
| **13** | **HSP90B1** | **76** |
| **13** | **MAP1LC3B** | **76** |
| **15** | **TUBA1B** | **74** |
| **16** | **SRRM1** | **62** |
| **17** | **ATP5B** | **60** |
| **18** | **TUBB4B** | **51** |
| **19** | **DNAJC7** | **50** |
| **19** | **SOD1** | **50** |
| **21** | **FXR1** | **49** |
| **22** | **NDUFV2** | **46** |
| **23** | **VDAC1** | **44** |
| **23** | **SRRM2** | **44** |
| **25** | **PTGES3** | **43** |

**Table 5:** Top 25 genes of the hippocampus region identified from the bottleneck centrality algorithm analysis.

| **Rank** | **Name** | **Score** |
| --- | --- | --- |
| **1** | **CUL3** | **1674** |
| **2** | **YWHAH** | **423** |
| **3** | **EP300** | **414** |
| **4** | **UBC** | **407** |
| **5** | **ILF3** | **302** |
| **6** | **KDM1A** | **105** |
| **7** | **NDUFS3** | **102** |
| **8** | **MAP1LC3B** | **91** |
| **9** | **RPN1** | **82** |
| **10** | **PPP2R1A** | **76** |
| **11** | **CSNK2B** | **69** |
| **12** | **COPS6** | **63** |
| **13** | **HDAC1** | **60** |
| **14** | **HSPA5** | **59** |
| **15** | **PHB** | **56** |
| **16** | **PSMC5** | **54** |
| **17** | **HSP90AB1** | **51** |
| **17** | **COX4I1** | **51** |
| **19** | **RBX1** | **50** |
| **20** | **TUBB** | **46** |
| **21** | **SMC3** | **45** |
| **22** | **SOD1** | **41** |
| **23** | **PSMA1** | **39** |
| **24** | **DNAJC7** | **38** |
| **25** | **PTGES3** | **35** |

### 3.4. Post-transcriptional regulation of the hub genes

Dysregulated expression of miRNA in AD has been reported by several lines of studies. Here, we have used the hub genes of **Table 2, 3, 4** and **5** to identify miRNAs that are known to regulate them and thereafter associate the findings with available literature about miRNAs that are differentially expressed in the diseased condition. For each of the hub genes, a set of miRNAs that are predicted to bind to the 3’ UTR region of the gene was obtained. Following processing in the R software platform there were 2540 and 2523 miRNA-transcript pair obtained from the entorhinal cortex and hippocampus region respectively. hsa-miR-3613-3p is seen to regulate the greatest number of hub genes, 18 in EC and 22 in hippocampus. In the entorhinal cortex this is followed closely by the hsa-miR-548 family and hsa-miR-4282, which regulate about 17 hub genes each and in the hippocampus by hsa-miR-7-1-3p, hsa-miR-7-2-3p, hsa-miR-325-3p, hsa-miR-6809-3p, hsa-miR-548 family which regulate ∼18 hub genes.

Kumar *et al*. 2017 in their study, reported differential expression of seven miRNAs, namely, hsa-miR-455-3p, hsa-miR-3613-3p, hsa-miR-4668-5p, hsa-miR-122-5p, hsa-miR-5001-5p, hsa-miR-4674 and hsa-miR-4741 in postmortem AD patients (Kumar, Vijayan and Reddy, 2017).In our study, we observed that the hub genes that we identified from PPI network analysis are also regulated by the seven differentially expressed miRNA. After filtering on the basis of context score percentile, setat 75, target genes that met the cutoff remained and are shown in (**Table 6 & 7)**. It can be seen from **Table 6 & 7**, few of the genes namely HSPA2 and NDRG1 in the entorhinal cortex and FXR1 in the hippocampus, highlighted in grey, show higher gene expression notwithstanding miRNA regulation, however, upon further analysis it was found that they are regulated by another set of miRNAs, hsa-miR-299 and hsa-miR-133a, which have been reported to be down-regulated by more than two-fold in AD (Zhang *et al*. 2016).

**Table 6:** List of miRNAs regulating hub genes, showing the number of target genes and the specific genes that are regulated by each miRNA from the entorhinal cortex. (* as reported by Kumar *et al*. 2017; **upregulated target genes are highlighted in grey)

| miRNA | Gene count | logFC* | Gene Symbol |
| --- | --- | --- | --- |
| hsa-miR-455-3p.1 | 4 | 11.3 | GPI (91), KPNA1(99), PSMD8(99) |
| hsa-miR-3613-3p | 6 | 3.67 | BRD4(80), GPI (83), HSPA2(92)** **, PSMB2(95), PSMB3(99), SKP1(86),** |
| hsa-miR-4668-5p | 5 | 3.38 | CDK5(94), PKM (85), PSMB2(90), SF3B5(91) |
| hsa-miR-122-5p | 3 | -3.85 | PKM (99), PRDX6(93) |
| hsa-miR-5001-5p | 6 | 2.37 | GPI (91), HSPBP1(82), PKM (91), POLR2L (93) |
| hsa-miR-4674 | 6 | 5.62 | KPNA1(80), OPTN (81), PIN1(86), PSMB2(83), RUVBL2 (95), SKP1(85) |
| hsa-miR-4741 | 4 | 2.07 | NDRG1(92) **\*\***, PKM (95), POLR2L (88), USP11(95) |

### 3.5. Convergent functional genomics analysis

The 13 genes found to be common between MCC and BN centrality analysis in the entorhinal cortex were queried in the Alzdata database to identify those genes that are expressed at the early stage of the disease and therefore likely to play a key role in disease pathogenesis. Of the 13 genes, BRD4, PPP1CB, USP11 and BAG3 have been found to be differentially expressed in early AD that is before the appearance of classical AD symptoms. BAG3, PSMD14, SKP1, BSG, HGS, KAT2B, IGF1R and PPP1CB also interact with core AD genes PSEN1, PSEN2, APP and MAPT. An early DEG, BRD4, is known to be a negative regulator of BACE1 while USP11 is an enzyme responsible for the removal of ubiquitin moiety from proteins. It is also known to be a positive regulator of HIF-1α pathway which when activated results in an increase in expression of glycolytic enzymes. In the entorhinal cortex, the pathway is likely inhibited due to the downregulation of USP11 leading to an overall reduction in the expression of glycolytic enzymes.

In the hippocampus region, of the 13 common genes that were used as an initial query, 3 genes, namely NDUFS3, RPN1 and SOD1 are early DEGs. 5 genes namely, HSP90AB1, KDM1A, PPP2R1A, SOD1 and PTGES3 interacts with core AD genes APP, PSEN1, PSEN2 and MAPT. Interestingly, PTGES3, a downregulated hub gene and also an interactor of core AD genes, is a negative regulator of HIF-1α through its interaction with PHD 1/2/3 regulatory proteins, which implies that unlike the entorhinal cortex where the HIF-1α pathway remains suppressed, the pathway remains active in the hippocampus and is likely to account for milder or less pronounced downregulation of glycolytic enzymes and glucose transporters as shown in **Table 8**.

## 4.0 Discussion

The study has identified 500 differentially expressed genes (DEGs) in the entorhinal cortex region that warrant special attention. Severe decline in gene expression of key glycolytic, pentose phosphate pathway and glycogen synthesis enzymes namely glucose phosphate isomerase (GPI), phospho-fructo kinase (PFK), triose phosphate isomerase 1 (TPI1), pyruvate kinase (PKM), glucose 6 phosphate dehydrogenase and phosphoglucomutase 1 (PGM1) has been observed. MCODE clustering analysis as well as bottleneck centrality analysis identified GPI, TPI, PKM and PGM1 hub genes potentially implicated in the disease pathogenesis. It is possible that the variation in gene expression for GPI and PKM could be because of upregulation of miRNA hsa-miR-455-3p.1, hsa-miR-3613-3p, hsa-miR-5001-5p and hsa-miR-4668-5p (**Table 6**). We also observed upregulation of PRDX6 in **Fig. 4(a)**, which from miRNA analysis, it seems is likely because of downregulation of miRNA hsa-miR-122-5p. Another striking observation from the entorhinal cortex is dysregulation of PIN1 gene expression. PIN1 knock-out mice develops AD pathology in an age-dependent manner. Also, loss of the prolyl-isomerase enzyme function has been reported to drive AD pathology through tau hyperphosphorylation mediated by GSK3, which has an affinity towards the cis-isoform of tau (Kondo *et al*. 2016). Additionally, PIN1 inhibits AMP-activated kinase (AMPK) activity thus inhibiting ATP-consuming anabolic processes and promoting catabolic processes that generate ATP (Herzig and Shaw, 2017). Even though we found no change in AMPK expression, PIN1 emerged as a severely downregulated hub gene. This implies lack of regulation of phosphorylation of ACC 1/2 (acetyl-CoA carboxylase 1/2) inhibits the formation of malonyl-CoA which is the substrate for fatty acid biosynthesis and an allosteric inhibitor of mitochondrial carnitine-palmitoyl transferase 1 enzyme. This indicates that in the later stage of the disease there is a global decline in anabolic processes. From our study, we have observed that PIN1 downregulation in the entorhinal cortex region is possibly mediated by the upregulation of miRNA hsa-miR-4674 (**Table 6**).

Downregulation of the global transcription factor, BRD4, a key hub gene in our study, is identified as an early DEG from CFG analysis. From post-transcriptional regulation analysis, it may be inferred that upregulation of hsa-miR-3613-3p leads to an observed decrease in expression of BRD4 (**Table 6**). Involvement of BRD4 with Aβ pathology has been reported by Zhang *et al*. 2022 through the inhibition of BRD4 on treatment by JQ1 and ARV-825 finally increasing the BACE1 activity by a post-transcriptional mechanism that remains to be studied (Zhang *et al*. 2022). Its downregulation therefore would subsequently lead to enhanced Aβ production, which likely represents an early event in Alzheimer’s disease (AD) pathogenesis.

Broadly speaking, autophagy is of three types, microautophagy, macroautophagy and chaperone mediated autophagy (Yang and Klionsky, 2009), all of which are implicated in the EC. HRS/HGS or hepatocyte growth factor receptor substrate is a component of the ESCRT-0 subcomplex and recruits the other three ESCRT subcomplexes on the endosomal surface (Wenzel *et al*. 2018). In our study, HGS is identified as a central hub gene in both MCC and BN centrality analysis and is severely downregulated suggesting an overall inhibition of microautophagy in late-stage AD. The process of macroautophagy proceeds with the formation of autophagosomes. Under stress conditions inhibition of PSMD14, interferes with the process of autophagosome biogenesis as it leads to the sequestration of the ATG9A, required for induction and nucleation of the phagophore, at the Golgi. In bulk microarray dataset, PSMD14, a central hub gene, is severely downregulated implying breakdown of macroautophagy process in late-stage AD. Aβ, a known inducer of oxidative stress, causes oxidative damage to proteins eventuating protein misfolding finally triggering the chaperone mediated autophagy (CMA) pathway, as indicated by the upregulation of BAG3 gene, a co-chaperone for HSP70 family of proteins involved in CMA (Gamerdinger *et al*. 2011).

Proteostatic stress resulting from accumulation of misfolded and otherwise non-functional proteins inside neurons is a well-studied feature of Alzheimer’s disease. In our study, in EC, out of three broadly classified autophagic pathways, only CMA has been observed to be functional. Since autophagy is a primary system for the removal and clearance of worn out, misfolded and aggregated proteins therefore dysregulation of key and accessory proteins of the micro and macro-autophagy pathway is likely a reason for proteostasis failure observed in late-stage AD.

In the hippocampus region, we observed, AD pathology, begin with excess production of Aβ, through enhanced BACE1 activity, driven by concomitant upregulation of FXR1 gene, which promotes BACE1 translation initiation, and downregulation of RTN3 gene, which is known to interact with BACE1 and negatively regulate its protease activity. Furthermore, the role of regulatory miRNAs is of great interest in the study of AD pathology. Of particular interest are hsa-miR-455-3p, hsa-miR-5001-5p, hsa-miR-4674 and hsa-miR-4741, since, from Targetscan database, it is observed that it regulates YWHAH (**Table 7**). YWHAH, a member of the 14-3-3 family of proteins, is known to be involved in diverse cellular processes. Of interest, is its role in autophagy-lysosomal (AL) pathway, where it is known to sequester phosphorylated-TEFB (p-TEFB), in the cytosol, preventing its nuclear translocation and hence expression of autophagy genes under nutrient rich conditions (Xu *et al*. 2019). Aggregation of misfolded proteins and oxidised proteins is a characteristic of AD possibly requiring continuous upregulation of the AL pathway keeping YWHAH expression down through miRNA mediated gene silencing initiated by hsa-miR-455-3p, hsa-miR-5001-5p, hsa-miR-4674 and hsa-miR-4741. However, it has a counterintuitive effect that exacerbates AD pathology since YWHAH has an important role in dopamine synthesis which acts both as a neurotransmitter and a catecholamine precursor of several important signalling molecules one of which is homovanillic acid. Morimoto *et al*. 2017 reported homovanillic acid as an early biomarker for AD and its CSF level decreases after the clinical onset of the disease (Morimoto et al., 2017). In our study, we observed that this could be due to downregulation of YWHAH gene expression as well as that of ALDH3A2 (−1.55) which catalyses the formation of homovanillic acid from dopamine.

**Table 7:**
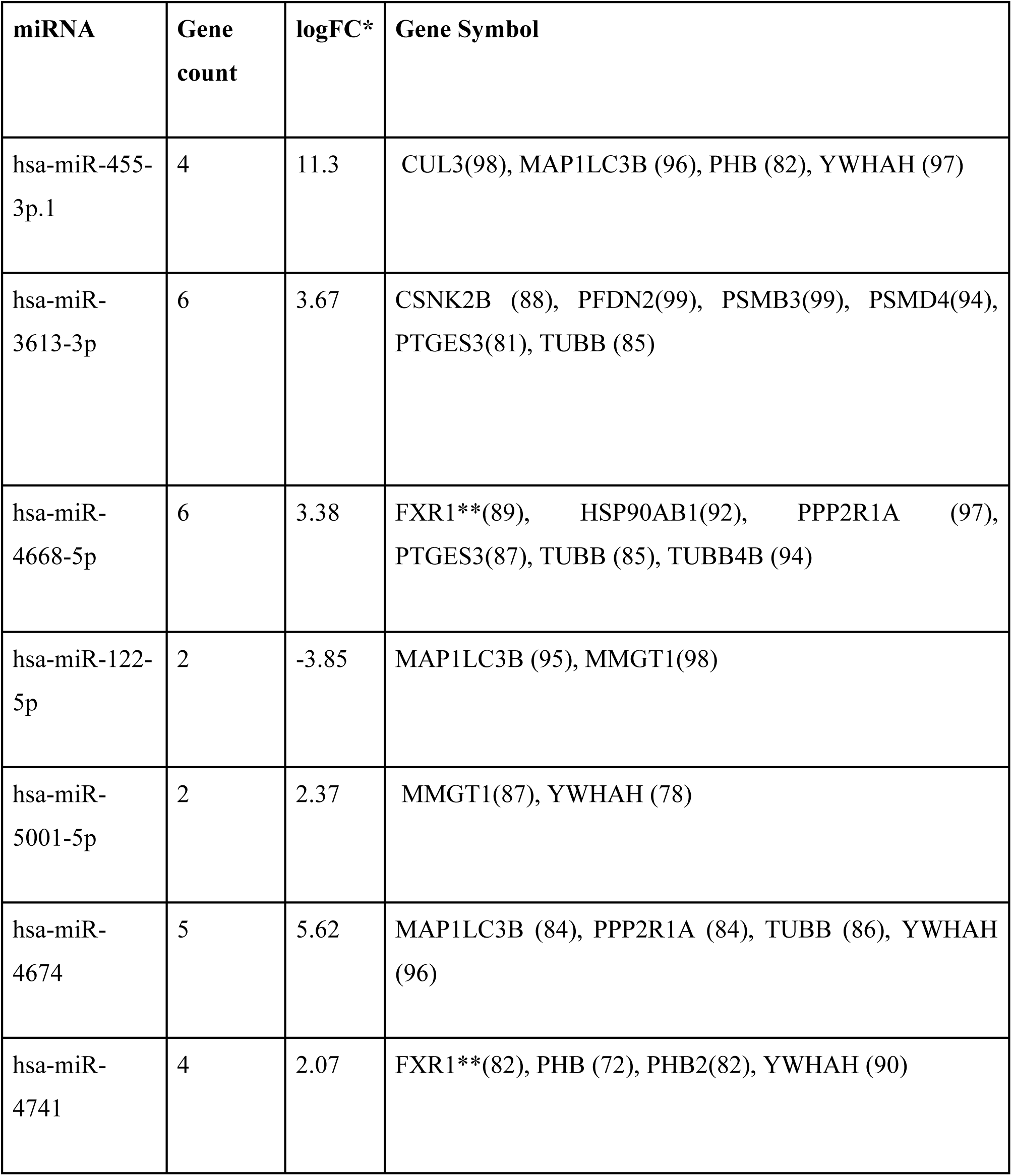
List of miRNAs regulating hub genes, showing the number of target genes and the specific genes that are regulated by each miRNA from the hippocampus. (* as reported by Kumar *et al*. 2017, **upregulated target genes are highlighted in grey)

| miRNA | Gene count | logFC* | Gene Symbol |
| --- | --- | --- | --- |
| hsa-miR-455-3p.1 | 4 | 11.3 | CUL3(98), MAP1LC3B (96), PHB (82), YWHAH (97) |
| hsa-miR-3613-3p | 6 | 3.67 | CSNK2B (88), PFDN2(99), PSMB3(99), PSMD4(94), PTGES3(81), TUBB (85) |
| hsa-miR-4668-5p | 6 | 3.38 | FXR1**(89), HSP90AB1(92), PPP2R1A (97), PTGES3(87), TUBB (85), TUBB4B (94) |
| hsa-miR-122-5p | 2 | -3.85 | MAP1LC3B (95), MMGT1(98) |
| hsa-miR-5001-5p | 2 | 2.37 | MMGT1(87), YWHAH (78) |
| hsa-miR-4674 | 5 | 5.62 | MAP1LC3B (84), PPP2R1A (84), TUBB (86), YWHAH (96) |
| hsa-miR-4741 | 4 | 2.07 | FXR1**(82), PHB (72), PHB2(82), YWHAH (90) |

**Table 8:** Change in gene expression in logFCof key glycolytic enzymes and glucose transporters in the entorhinal cortex and the hippocampus.

| Gene Symbol | LogFC entorhinal cortex | LogFC hippocampus |
| --- | --- | --- |
| GPI | -2.84 | -1.96 |
| TPI | -1.76 | -1.11 |
| SLC2A6 | -1.26 | 0.66 |
| PFKM | -2.05 | -1.53 |
| PKM | -1.41 | -0.70 |

In conclusion, our study reports involvement of distinct cellular pathways driving tau and amyloidogenic pathogenesis in the entorhinal cortex and hippocampus from microarray data analysis. In the EC, increase in BACE1 activity through concomitant miRNA mediated downregulation of BRD4 and upregulation of the Oncostatin-M-pathway leads to enhanced amyloidogenic processing and Aβ production. Interestingly, the same effect in the hippocampus is driven through the differential expression of FXR1 and RTN3 gene. Thus, though BACE1 is central in promoting accumulation of toxic Aβ peptides in both the region however the molecular mechanism of its regulation is distinct shedding light on the fact that it is perhaps not possible to develop AD therapeutics using single target.

## Data availability statement

Data used in the study have been retrieved from NCBI repository GEO DataSets (GSE5281) avail-able at https://www.ncbi.nlm.nih.gov/geo/query/acc.cgi?acc=GSE5281.

## Declaration

We declare no competing interest.

## Acknowledgements

The authors acknowledgeThe Centre for High Performance Computing in Modern Biology, University of Calcutta, for providing every support required for this work. Moumita Biswas acknowledges the University Grants Commission for her fellowship (695/CSIRNETJRF2019).

