## Supplementary material for "DIFFERENTIAL REGULATION OF GENES IN ENTORHINAL CORTEX AND HIPPOCAMPUS IN LATE ONSET AD": Table S1

| **Gene.Symbol** | **Gene.Title** | **logFC** | **adj.P.Val** |
| --- | --- | --- | --- |
| **FAM107B** | **family with sequence similarity 107, member B** | **3.202855034** | **2.97E-07** |
| **TJP2** | **tight junction protein 2** | **3.012337349** | **2.43E-05** |
| **HIPK2** | **homeodomain interacting protein kinase 2** | **2.985201764** | **2.62E-05** |
| **SPP1** | **secreted phosphoprotein 1** | **2.857486027** | **9.66E-05** |
| **ERBB2IP** | **erbb2 interacting protein** | **2.830122073** | **1.48E-05** |
| **C21orf91** | **chromosome 21 open reading frame 91** | **2.648931515** | **6.61E-06** |
| **SLCO1A2** | **solute carrier organic anion transporter family, member 1A2** | **2.614775316** | **1.90E-06** |
| **HSPA2** | **heat shock 70kDa protein 2** | **2.532600743** | **1.52E-05** |
| **MAFF** | **v-maf avian musculoaponeurotic fibrosarcoma oncogene homolog F** | **2.427045883** | **8.89E-05** |
| **MOBP** | **myelin-associated oligodendrocyte basic protein** | **2.402481348** | **2.21E-06** |
| **GOLIM4** | **golgi integral membrane protein 4** | **2.386644276** | **1.85E-05** |
| **IRF2BP2** | **interferon regulatory factor 2 binding protein 2** | **2.347295671** | **1.31E-05** |
| **PMP2** | **peripheral myelin protein 2** | **2.343488616** | **0.000141034** |
| **CD44** | **CD44 molecule (Indian blood group)** | **2.30910521** | **0.00013854** |
| **PALLD** | **palladin, cytoskeletal associated protein** | **2.30394558** | **0.000147256** |
| **RFX4** | **regulatory factor X, 4 (influences HLA class II expression)** | **2.298214038** | **7.63E-05** |
| **CLDN11** | **claudin 11** | **2.276636151** | **1.39E-05** |
| **KAT2B** | **K(lysine) acetyltransferase 2B** | **2.276458687** | **0.000782899** |
| **ZIC1** | **Zic family member 1** | **2.211227266** | **2.02E-05** |

**Table S1: Top 20 up-regulated genes in the entorhinal cortex of the dataset GSE5281**
