## Supplementary material for "DIFFERENTIAL REGULATION OF GENES IN ENTORHINAL CORTEX AND HIPPOCAMPUS IN LATE ONSET AD": Table S2

| **Gene.Symbol** | **Gene.Title** | **logFC** | **adj.P.Val** |
| --- | --- | --- | --- |
| **MIF** | **macrophage migration inhibitory factor (glycosylation-inhibiting factor)** | **-3.674762961** | **2.58E-06** |
| **CHRM1** | **cholinergic receptor, muscarinic 1** | **-3.436151201** | **6.37E-06** |
| **STYK1** | **serine/threonine/tyrosine kinase 1** | **-3.366952538** | **3.20E-05** |
| **RBP4** | **retinol binding protein 4, plasma** | **-3.228314184** | **2.34E-05** |
| **AP2M1** | **adaptor-related protein complex 2, mu 1 subunit** | **-3.201108558** | **2.65E-05** |
| **PRDX2** | **peroxiredoxin 2** | **-3.097893275** | **2.58E-06** |
| **TECR** | **trans-2,3-enoyl-CoA reductase** | **-2.951245365** | **1.94E-06** |
| **VPS28** | **vacuolar protein sorting 28 homolog (S. cerevisiae)** | **-2.917047526** | **2.21E-06** |
| **TUBB4A** | **tubulin, beta 4A class IVa** | **-2.849997664** | **1.31E-05** |
| **GPI** | **glucose-6-phosphate isomerase** | **-2.842224274** | **2.75E-05** |
| **YJEFN3** | **YjeF N-terminal domain containing 3** | **-2.833898632** | **4.22E-06** |
| **GPR88** | **G protein-coupled receptor 88** | **-2.779575612** | **3.03E-05** |
| **LIN37** | **lin-37 homolog (C. elegans)** | **-2.744599648** | **3.21E-06** |
| **RNF145** | **ring finger protein 145** | **-2.730962752** | **8.53E-05** |
| **ATP5D** | **ATP synthase, H+ transporting, mitochondrial F1 complex, delta subunit** | **-2.717413108** | **7.10E-06** |
| **C1QTNF4** | **C1q and tumour necrosis factor related protein 4** | **-2.69267969** | **0.000100611** |
| **PCSK1N** | **proprotein convertase subtilisin/kexin type 1 inhibitor** | **-2.62116477** | **1.10E-05** |
| **ALKBH6** | **alkB, alkylation repair homolog 6 (E. coli)** | **-2.599163118** | **1.63E-06** |
| **CALY** | **calcyon neuron-specific vesicular protein** | **-2.596116204** | **0.00014125** |

**Table S2: Top 20 down-regulated genes in the entorhinal cortex of the dataset GSE5281**
