## Supplementary material for "DIFFERENTIAL REGULATION OF GENES IN ENTORHINAL CORTEX AND HIPPOCAMPUS IN LATE ONSET AD": Table S3

| **Gene.Symbol** | **Gene.Title** | **logFC** | **Adj.P.Val** |
| --- | --- | --- | --- |
| **LOC202181** | **SUMO-interacting motifs containing 1 pseudogene** | **2.566865** | **0.000481** |
| **FXR1** | **fragile X mental retardation, autosomal homolog 1** | **2.447927** | **1.01E-05** |
| **MINK1** | **misshapen-like kinase 1** | **2.392499** | **3.33E-06** |
| **SRRM2** | **serine/arginine repetitive matrix 2** | **2.384078** | **4.78E-05** |
| **NOVA2** | **neuro-oncological ventral antigen 2** | **2.351362** | **5.20E-09** |
| **HRK** | **harakiri, BCL2 interacting protein** | **2.314022** | **3.08E-05** |
| **RNF165** | **ring finger protein 165** | **2.281456** | **2.69E-05** |
| **SMC3** | **structural maintenance of chromosomes 3** | **2.253756** | **3.13E-05** |
| **TTBK2** | **tau tubulin kinase 2** | **2.244158** | **1.31E-06** |
| **HSP90B1** | **heat shock protein 90kDa beta (Grp94), member 1** | **2.243864** | **1.77E-05** |
| **BBX** | **bobby sox homolog (Drosophila)** | **2.146831** | **9.13E-07** |
| **GGA3** | **golgi-associated, gamma adaptin ear containing, ARF binding protein 3** | **2.109596** | **1.23E-10** |
| **PDCD6** | **programmed cell death 6** | **2.06771** | **0.000914** |
| **IQCA1** | **IQ motif containing with AAA domain 1** | **2.028709** | **0.000656** |
| **SEMA6A** | **sema domain, transmembrane domain (TM), and cytoplasmic domain, (semaphorin) 6A** | **1.975249** | **3.56E-05** |
| **NSUN6** | **NOP2/Sun domain family, member 6** | **1.967551** | **0.000452** |
| **ANKRD13D** | **ankyrin repeat domain 13 family, member D** | **1.960319** | **4.40E-05** |
| **SEPT7P2** | **septin 7 pseudogene 2** | **1.91218** | **5.21E-05** |
| **COL27A1** | **collagen, type XXVII, alpha 1** | **1.889789** | **0.000384** |
| **RNPC3** | **RNA-binding region (RNP1, RRM) containing 3** | **1.880661** | **0.000193** |

**Table S3: Top 20 up-regulated genes in the hippocampus region of the dataset GSE5281**
