## Supplementary material for "DIFFERENTIAL REGULATION OF GENES IN ENTORHINAL CORTEX AND HIPPOCAMPUS IN LATE ONSET AD": Table S4

| **Gene.Symbol** | **Gene.Title** | **logFC** | **Adj.P.Val** |
| --- | --- | --- | --- |
| **YWHAH** | **tyrosine 3-monooxygenase/tryptophan 5-monooxygenase activation protein, eta** | **-3.51784** | **7.47E-05** |
| **ARPC1A** | **actin related protein 2/3 complex, subunit 1A, 41kDa** | **-2.93622** | **0.000451** |
| **CASD1** | **CAS1 domain containing 1** | **-2.60892** | **0.000132** |
| **CD9** | **CD9 molecule** | **-2.54875** | **0.000157** |
| **RTN3** | **reticulon 3** | **-2.52527** | **9.33E-07** |
| **HSP90AB1** | **heat shock protein 90kDa alpha (cytosolic), class B member 1** | **-2.49007** | **0.00015** |
| **PDGFA** | **platelet-derived growth factor alpha polypeptide** | **-2.40952** | **1.31E-06** |
| **NDUFA10** | **NADH dehydrogenase (ubiquinone) 1 alpha subcomplex, 10, 42kDa** | **-2.3948** | **1.47E-06** |
| **DHCR24** | **24-dehydrocholesterol reductase** | **-2.37005** | **0.000216** |
| **MMADHC** | **methylmalonic aciduria (cobalamin deficiency) cblD type, with homocystinuria** | **-2.34316** | **0.000953** |
| **IDH3A** | **isocitrate dehydrogenase 3 (NAD+) alpha** | **-2.31939** | **0.000655** |
| **TTC19** | **tetratricopeptide repeat domain 19** | **-2.30693** | **2.39E-05** |
| **MTURN** | **maturin, neural progenitor differentiation regulator homolog (Xenopus)** | **-2.30302** | **0.000127** |
| **TUBB** | **tubulin, beta class I** | **-2.27094** | **2.87E-05** |
| **GOT1** | **glutamic-oxaloacetic transaminase 1, soluble** | **-2.26829** | **0.000708** |
| **ATP6V1H** | **ATPase, H+ transporting, lysosomal 50/57kDa, V1 subunit H** | **-2.24445** | **0.000485** |
| **PSMB3** | **proteasome (prosome, macropain) subunit, beta type, 3** | **-2.23251** | **5.16E-05** |
| **SCN2B** | **sodium channel, voltage-gated, type II, beta subunit** | **-2.23035** | **5.04E-06** |
| **EAPP** | **E2F-associated phosphoprotein** | **-2.22571** | **2.53E-05** |
| **ADAM23** | **ADAM metallopeptidase domain 23** | **-2.21122** | **0.000111** |

**Table S4: Top 20 down-regulated genes in the hippocampus region of the dataset GSE5281**
